# A new simulation framework to evaluate the suitability of eDNA for marine and aquatic Environmental Impact Assessments

**DOI:** 10.1101/2022.12.15.520594

**Authors:** J. Coston-Guarini, S. Hinz, L. Mirimin, J.-M. Guarini

## Abstract

This study evaluates how eDNA information could be used within Environmental Impact Assessment (EIA). We developed an original model to simulate the conditions for which an eDNA signal detects, or does not detect, an impact on a targeted (receptor) species in a given project area. The simulation has four consecutive steps. First, a deterministic model simulated the dynamics of the receptor population and their eDNA fragment concentrations in the environment. Second, random distributions of receptor organisms and eDNA fragment quantities at steady-state were simulated within the project area. Then Simple Random Samplings were performed for both the receptor and eDNA. Third, post-sampling processes (eDNA extraction, amplification, analysis) were simulated to estimate the detection probability of the species from sample plan characteristics (size of sampling unit, number of samples collected). Fourth, we simulated an impact by modifying the growth, mortality and mobility (null, passive and active) parameters of the receptor species, then determined if an impact was detected. Detection probability curves were estimated for a range of sample volumes fitted with a Weibull cumulative distribution function. An F-like statistic compared detection curves before and after impact. Twelve scenarios were simulated. A statistically significant impact was detected with eDNA when receptor species growth rate was halved, but only in cases of null or passive mobility. When the receptor experienced both reduced growth and increased mortality rates, an impact was detected in all three mobility cases (null, passive and active). Our results suggest that an impact could be detected using eDNA if both the population dynamics of the receptor and the dynamics of DNA shed into the environment are known. These results indicate that caution should be exercised with eDNA data for EIA, but the proposed framework provides a valuable starting point to improve interpretation of indirect observation methods such as eDNA.

## 1 Introduction

The increasing capacity to sequence and identify species from DNA fragments captured within environmental samples has fueled an ambition to see DNA analyses integrated within the environmental sciences (Lodge et al., 2012; Kelly et al., 2014; Deiner et al., 2017). Studies and applications in conservation that had previously relied primarily on visual observation of organisms’ presence are now exploring how to complement or even replace observation with environmental DNA (*e*.*g*. Ruppert et al. 2019). Because genetic material, ‘environmental DNA’ (eDNA) is being extracted from any type of environmental matrix, (*e*.*g*. water, soils, sediments, other organisms, and atmospheric samples), recently, a new potential area of application - Environmental Impact Assessments - has been suggested Hinz et al. (2022).

Environmental Impact Assessment (EIA) is a formal process for assessing the impact(s) or effects of different human activities on their surrounding environments. It was adopted as a policy tool beginning in the early 1970s, and has since become a part of decision-making systems in a majority of countries (Wilson et al., 2017; UN Environment, 2018).Unfortunately, there is no universal definition of EIA, but the International Association for Impact Assessment has defined it as “[…*] the process of identifying the future consequences of a current or proposed action*”. In general, it is structured into five steps (called screening, scoping, baseline, impact assessment and monitoring) during which impacts are identified clearly so that necessary mitigation actions can be taken. In all cases, impact assessments are performed prior to the implementation of the project. Since EIA is a forecasting process (Coston-Guarini et al., 2017) where modeled estimates of impact will depend on the quality of background information available about the ecological system under evaluation, eDNA analyses would offer many advantages over traditional biodiversity survey methods used. In marine environments, where information on the composition of biological communities may be both outdated and insufficiently described with respect to the framework of impact assessments, there has been strong interest in adopting the technique (Kelly et al., 2014; Hinz et al., 2022).

The eDNA community is heavily invested in improving the quality and comparability of assays (*e*.*g*. Thalinger et al. 2021), and the precision and sensitivity of eDNA offers the possibility to detect and identify organisms’ DNA within most environmental matrices. However, ambiguities regarding the sources and fate of DNA are challenging within EIA. This is because DNA in an environmental sample can be from many different sources: intact cells of a living organism (such as phytoplankton or bacteria) and/or cells shed by an organism (*e*.*g*. saliva, mucus, skin), or from a dead organism. This diversity has inspired a debate in the literature about what ‘eDNA’ encompasses (*e*.*g*. Pawlowski et al. 2020; Rodriguez-Ezpeleta et al. 2021). These ambiguities complicate the re-use and interpretation of the data within ecological models because numerical tools require the source to be explicit. For instance, this requirement is needed for treating mixing, dispersion and transport. In addition, for EIA in particular, the statistical design of sampling programs that guide the collection of genetic material samples are important to being able to characterize patterns for evaluating a potential impact. One means to explore these issues is by considering environmental DNA within a simulation framework, using existing knowledge about ecological systems and eDNA.

The approach used for impact assessments today, originated in statistical decision theory (Leopold et al., 1971). This means being able to decide if a difference, *i*.*e*. a change in an estimator (in this case the eDNA signal) is significant or not (Hinz et al., 2022). In practice, a comparison is made between values measured before and after a project begins, and both within and outside of the strict bounds of the project area. The impact assessment defines the conditions for which an impact can be detected in an ecosystem (Coston-Guarini et al., 2017). To do this, a set of “receptors” for the potential impact are defined. Receptors are objects (organisms, habitats, communities, …) expected to have a detectable response (*e*.*g*. change in abundance, behaviour, distribution, …) when exposed to a disturbance. The objects selected are often subject to some type of existing regulation (such as protected species) or are of special interest to a project area. Thus, the potential impact is assessed for receptors within the strict geographic bounds of the project area and relative to the surrounding area. When inserted in a regulatory framework, changes in receptors can lead to legal issues if threshold values are crossed; this makes a statistical assessment politically sensitive. This may concern: endangered species, sensitive habitats, and harmful and pathogenic species. A monitoring survey may also be performed *a posteriori* after project implementation, to ensure that mitigation measures taken to decrease any potential impacts are indeed effective (Wilson et al., 2017).

Despite a handful of preliminary initiatives (*e*.*g*. Pochon et al. 2015; Decher et al. 2016; Cordier et al. 2017; Laroche et al. 2017; Stoeck et al. 2018; Cordier et al. 2019; Forster et al. 2019; Brandt et al. 2020; Jonsson et al. 2020; Lanzen et al. 2021; Lejzerowicz et al. 2021; Mauffrey et al. 2021) few of these have dealt explicitly with the use of eDNA within a formal EIA process. And, while some reactive-transport modelling of DNA molecules has been performed (*e*.*g*. Andruszkiewicz et al. 2019), none of this work can be extended within EIA. Therefore we have developed a numerical forecasting tool that evaluates the suitability of eDNA for environmental impact assessment. The model presented in this article simulates *in silico* a generic impact on an aquatic (or marine) ecosystem, and performs both direct estimates for receptors and indirect estimates for the DNA fragments released by these receptors into the surrounding environment. The framework includes the eDNA analytical process (inspired by Furlan et al. (2016)) and then estimates probabilities of detection. Finally, the results of both the organism-based and eDNA-based detection of the receptor species in the ecosystem are considered within a statistical impact assessment framework (Coston-Guarini et al., 2017).

## 2 Model conception

The simulation framework described in this article is designed to compare impact scenarios using either a traditional organism sampling approach or a set of environmental DNA analyses. The general definition of eDNA proposed “total pool of DNA isolated from environmental samples” cited in Pawlowski et al. (2020) cannot be used in an ecosystem model. This is because an ecosystem model requires a mass balance of the objects (*e*.*g*. organisms, molecules) represented in the entire ecosystem, not just in the sample. The model definition - *the total pool of DNA shed by organisms into the ecosystem, whether they are alive or dead* - is based on the discussion in (Barnes & Turner, 2015). The model quantifies changes in a single receptor species, designated during the screening and scoping phases of an EIA, for an aquatic system (Figure 1). Project implementation is presumed to affect directly the receptor species dynamics by modifying the ecological performance of individuals. Hence, impact is defined as any process that would affect the ecophysiology of the receptor, by reducing growth rate and/or increasing mortality rate. This is a prolongation of an earlier model of EIA (Coston-Guarini et al., 2017), where a framework was designed to simulate the dynamics of a receptor population, quantifying the population states within a project area and in connection with the surrounding environment.

**Figure 1:**
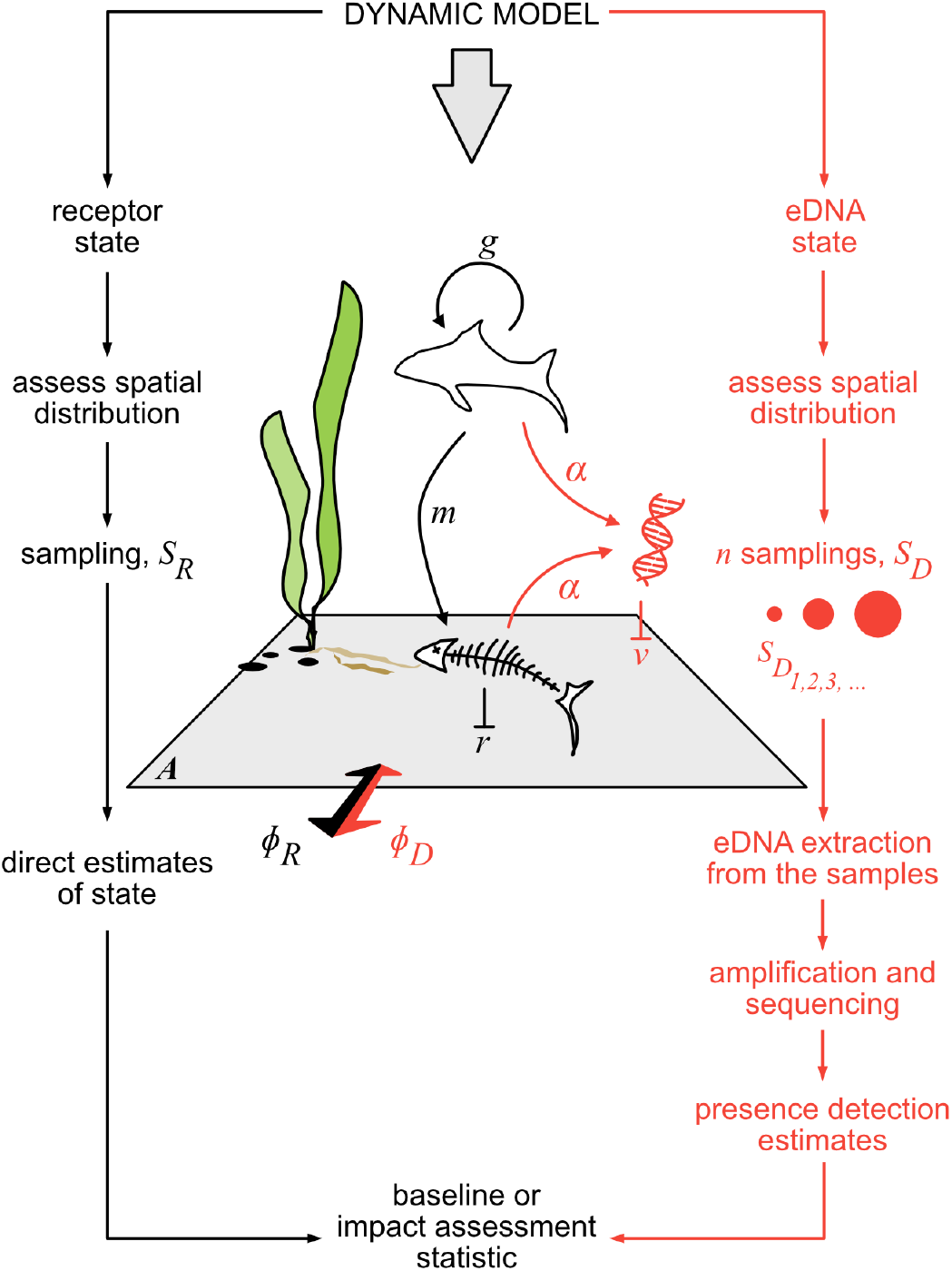
Conceptual key to the model presented in this article. A receptor species (R) and its DNA (D) are treated with either traditional methods (*S*_*R*_, left-hand side with black text and arrows) or by quantification of their environmental DNA signal (*S*_*D*_, right-hand side with orange text and arrows). A dynamic model links the population of the receptor species (growth (*g*), mortality (*m*), recycling (*r*)) with their DNA shed at a rate of *α* into the environment (*ν* is the rate of decay). Impact is always estimated relative to a project area (gray trapezoid, *A*) and both individual organisms and their DNA can exchange, *ϕ*, with the surrounding environment.

Figure 1 illustrates the two simulation pathways developed in the next sections. On the left side (in black text), a receptor species is treated with traditional information about the population presence. In this article, we have chosen not to discuss the population dynamics of a particular species. The model was designed as a demonstration tool, that aggregates the current state of knowledge without loss of generality. On the right side, in red, is the succession of steps for treating the same species with its eDNA signal, only. The DNA pool was considered to have multiple potential sources contributing to the signal of the same species and the DNA shed was assumed to be both passively and isotropically dispersed. First, the shedding dynamics of eDNA released by both living and dead organisms of the receptor species (section 3.2) are estimated. This also could allow for the possible contribution from resuspension of sedimented DNA (*e*.*g*. Sakata et al. 2020); however this is not explicitly included in the present version of the model. Then the spatial distribution of the state variables within the project area are treated, and any sampling issues for the receptor and DNA fragments released in the environment are addressed (section 3.3). Finally the impact statistic is forecast using a comparison with the receptor baseline of each approach (sections 3.4 and 3.5).

This model architecture permits the first quantified examination of how different aspects of the environmental conditions, sampling plan and laboratory and analytical pipelines interact to improve or reduce the interpretability of eDNA data. The model focuses on tracking the mass of DNA as it moves through the system because this is what determines the detectability of the species from an environmental sample. During development, we noted two areas of eDNA studies where large uncertainties limit a purely theoretical consideration: 1. sample sizes and 2. shedding rate dynamics. Shedding rate dynamics are parameterized using an allometric function because not enough systematic information is available for different taxonomic groups. In most eDNA studies, sample size appears to be chosen mostly as a result of trade-offs between practical sample processing aspects (e.g. amount of water prior to filter clogging and/or time and/or cost constraints; Tsuji et al. (2019); Allan et al. (2021)); the effects of sub-optimal sampling are often not considered fully (e.g. Buxton et al. (2021), see also supplementary information section A.6).

## 3 Model framework

### 3.1 Receptor submodel

The receptor population was defined by two state variables *R*_*I*_ and *M*_*I*_. *R*_*I*_ is the total abundance of living organisms of the receptor species assumed to be impacted within the project area. The corresponding species abundance of living organisms outside of the project area is represented as *R*_*E*_. The project area is considered as open, hence exchanges are allowed between *R*_*I*_ and *R*_*E*_. The number of dead receptor organisms within the project area is *M*_*I*_. A generic minimal formulation describes the dynamics of *R*_*I*_ and *M*_*I*_:

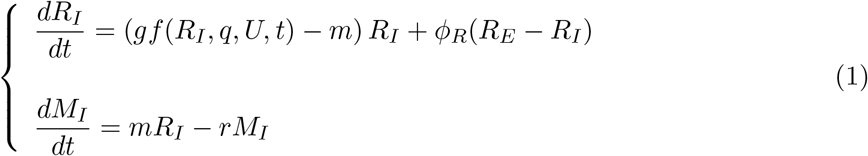

where *g* is a population growth rate (*time*^−1^), *f* is a growth limitation function (dimensionless) that depends on *R*_*I*_, a vector of parameters *q*, a vector of forcing factors, *U* and time *t, m* is a mortality rate (*time*^−1^), *ϕ*_*R*_ is a dispersion rate (*time*^−1^) and *r* is the recycling rate of dead organisms (*time*^−1^). It is assumed that at the scale of the dynamics, dead organisms remain within the project area, without outside exchanges.

A minimal formulation for the limitation function was then introduced:

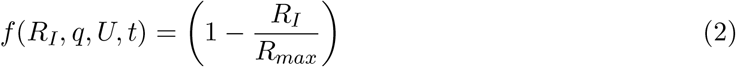

where *R*_*max*_ is the maximum quantity of *R*_*I*_ that the project area can sustain.

Next we formulated a generic case configured with a plausible set of parameter values (see Table 1). For the receptor population parameters, general allometric functions are used that apply to a large range of organism sizes (Peters, 1983). The sizes of individuals are defined as *L*, a linear dimension (unit of Length) of the body. The weight of the individuals (unit of Mass) is calculated as *W* = *a*_*L*_*L*^3^. The maximum density is then : *R*_*max*_ = *a*_*R*_*W* ^−1.0^. The growth and mortality rates are given by: *g* = *a*_*g*_*W* ^−0.25^ and *m* = *a*_*m*_*W* ^−0.25^. In addition, the individual weight, *W*, was used to convert abundances into the biomass value used by the eDNA submodel in the next section.

**Table 1.** Parameter values used for the impact system simulation discussed in this article.

| Parameter | Value (range) | Unit | Source |
| --- | --- | --- | --- |
| $A$, project area | $1 \times 10^5$ | $m^2$ | |
| $\rho$, scaling factor | 3.0 | $m$ | |
| $S_R$, sampling unit, receptor | 1.0 | $m^2$ | |
| $S_D$, sampling unit, eDNA | 0.004 - 0.060 | $m^3$ | |
| $L$, individual size | 0.050 | $m$ | |
| $a_L$, allometric coefficient, individual size | 0.0025 | $g.m^{-3}$ | [1] |
| $a_R$, allometric coefficient, maximum density | 10.0 | $nb \text{ of } ind.g^{-1}$ | [1] |
| $a_g$, allometric coefficient, individual growth | 0.1 | $day^{-1}$ | [1] |
| $a_m$, allometric coefficient, individual mortality | 0.003 | $day^{-1}$ | [1] |
| $\phi_R$, dispersion rate, receptor | 0.00 or 0.01 or 0.10 | $day^{-1}$ | |
| $\phi_D$, dispersion rate, eDNA | 0.01 | $day^{-1}$ | |
| $\alpha_W$, shedding rate | 81.6 (16.8 - 165.6) | $nb \text{ of } fragments.g^{-1}.day^{-1}$ | [2] |
| $\nu$, decay rate of eDNA | 0.771 (0.232 - 1.272) | $day^{-1}$ | [2] |
| $V_a$, aliquot volume | 5.0 | $ml$ | |
| $V_e$, elutant volume | 0.005 | $ml$ | |
| $x_c$, number of DNA fragments per aliquot volume specified by analytical protocol | 30.0 | $nb \text{ of } fragments.aliquot^{-1}$ | [3] |
| $b_x$, Weibull shape parameter | 7.0 | dimensionless | [3] |
| $p_{10}$, probability false positive | 0.1 | dimensionless | |
| $k$, number of replicate aliquots | 8 | *number* | |

Notes: [1] Peters (1983), [2] Sansom & Sassoubre (2017), [3] Furlan et al. (2016). Detailed justifications for all other values are presented in the supplementary section (Appendix A).

### 3.2 eDNA submodel

The fraction of eDNA shed by receptor species *R*_*I*_ (inside the project area) is defined as *D*_*I*_. The dynamics of *D*_*I*_ was formulated as:

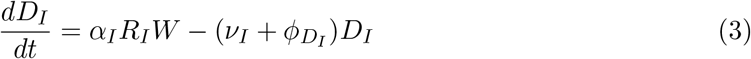

where *α*_*I*_ is an eDNA shedding rate (from living organisms) in *unit*_*D*_.*unit*_*RW*_ ^−1^.*time*^−1^, *ν*_*I*_, in *time*^−1^, an eDNA decay rate and *ϕ*_*D*_, in *time*^−1^, a dispersive rate of eDNA from within the project area to the surrounding area outside.

Even if *D*_*I*_ is defined as having been produced within the project area, it does not necessarily represent the quantity of eDNA in the project area for that same species. The *D*_*I*_ may be mixed with DNA coming from outside the project area and DNA shed from dead individuals within the project area. Therefore, we have set the eDNA quantity coming from a fraction of the species population which does not correspond to the receptor as *D*_*N*_ (because it is outside the project area), and its dynamics are described by:

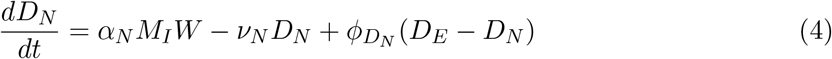

where *α*_*N*_ is a eDNA shedding rate in *unit*_*D*_.*unit*_*RW*_ ^−1^.*time*^−1^, *ν*_*N*_, in *time*^−1^, an eDNA loss term, *ϕ*_*D_N_*_ in *time*^−1^ is the exchange rate from outside to inside the project area, and *D*_*E*_ in *unit*_*D*_, is a quantity of eDNA outside of the project area.

Finally, the observed eDNA in the project area was defined as *D*, and its dynamics are formulated as:

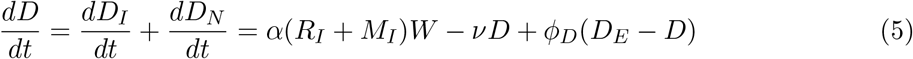

assuming that *α*_*I*_ = *α*_*N*_ = *α, ν*_*I*_ = *ν*_*N*_ = *ν* and 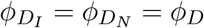. In this configuration, *D* can be reconstructed by assuming that the state of eDNA fragments is homogeneous over the distribution area, and only determined by a set of identical parameters, the shedding rate *α*, the decay rate *ν* and the exchange rate *ϕ*_*D*_.

The model defined by Equations 1, 2 and 5 is described by three state variables {*R*_*I*_, *M*_*I*_, *D*} with initial conditions {*R*_*I*_(0), *M*_*I*_(0), *D*(0)}, for which only *R*_*I*_ must be greater than 0. The processes governing the dynamics of the state variables include a simplified vector of seven parameters to estimate ({*g, R*_*max*_, *m, ϕ*_*R*_, *α, ν, ϕ*_*D*_}) and two forcing variables, *R*_*E*_ and *D*_*E*_, which represent the quantities of the receptor species and eDNA outside of the project area, respectively.

To complete the parameterization, shedding and decay rates were determined from values in the literature; however, the shedding rate *α* was expressed as *α*_*W*_ *W* where *α*_*W*_ is a number of molecules released per gram (of individual) per day (*nb*.*g*^−1^.*day*^−1^). This expression complies with the allometric parameterization of the receptor submodel. The decay rate is in *day*^−1^.

The exchange rate, *ϕ*_*D*_, in *day*^−1^, is a passive dispersion rate and is used as a reference for the model. This implies that, if *ϕ*_*R*_ = *ϕ*_*D*_, then the dispersion of the receptor is passive as well.

### 3.3 Sampling characteristics

Considering that the entire project area *A* is too large to be investigated, sampling plans are assumed to be implemented to estimate the state variables at a time *t*. The sampling plan designs must ensure that the estimators are not biased, and that they minimize as much as possible their uncertainty. This therefore requires an understanding of how the state variables are spatially distributed over the project area.

With respect to the receptor species, it is assumed that *R*_*I*_ is randomly spatially distributed over the project area *A*. Sampling design begins with the definition of a sampling plan, the sampling unit, *S*_*R*_, and the sample size *n*. The optimal sampling plan in this configuration is a Simple Random Sampling (SRS). In each sample *i, i* = {1, *n*}, the quantity of the receptor, *r*_*i*_ is determined, and *r*_*i*_ follows a Poisson distribution *r*_*i*_ ∼ *Pois*(*λ*_*R*_), where 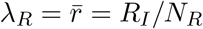, and *N*_*R*_ is the total number of sampling units than can be collected in the project area, *i*.*e. N*_*R*_ = *A/S*_*R*_. The variance of the *r*_*i*_ (*s*^2^) is of the same order of magnitude as the average 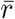. To ensure that the Poisson distribution can be approximated by a normal distribution, *S*_*R*_ was chosen in such a way that 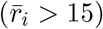. As *R*_*I*_(*t*) is calculated and *A* is fixed, *S*_*R*_ can be determined prior to sampling. In a case where *S*_*R*_ needs to be determined without knowing *R*_*I*_, a pre-sampling estimate must be performed.

Regarding a sampling plan for eDNA, the sampling unit *S*_*D*_ and the total number of samples, *m*, can differ significantly from the receptor. The spatial distribution of eDNA is random, with a possibility of exploring clumping effects at small scale. The optimal sampling plan is also a Random Simple Sampling (SRS). *S*_*D*_ is defined as a volume of the environment containing eDNA. The sample size (*i*.*e*. the number of samples taken), *m* is defined among *N*_*D*_, the total number of sampling units than can be collected, *i*.*e. N*_*D*_ = *ρA/S*_*D*_, where *ρ* is a scaling factor, to account for the dimension of the the environment in which eDNA is sampled (*i*.*e*. the area is converted to a volume). In each sample *j*, {*j* = 1, *m*}, when the spatial distribution is assumed to be purely random, the quantity of eDNA (in term of number of molecules) *d*_*j*_ follows a Poisson distribution *d*_*j*_ ∼ *Pois*(*λ*_*D*_), where 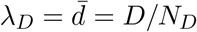. The variance of the sample set 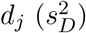 is of the same order of magnitude as the average 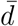. When the spatial distribution is assumed to exhibit aggregation, *d*_*j*_ follows a negative binomial distribution, *d*_*j*_ ∼ *B*_*neg*_(*r*_*B*_, *p*_*B*_), where *p*_*B*_ is a probability parameter, and *r*_*B*_ a dimension parameter equal to *r*_*B*_ = *λ*_*D*_(1 − *p*_*B*_)*/p*_*B*_. The more *r*_*B*_ increases, the more the distribution converges to a Poisson distribution with parameter *λ*_*D*_. The more *r*_*B*_ decreases, the more the DNA molecules are aggregated. There is no criteria to optimize *S*_*D*_.

### 3.4 Impact Assessment *per se*

In the sampling context defined in the previous section, an impact assessment is performed on the receptor by measuring the difference between two estimates of *R*_*I*_(*t*), before and after project implementation. *R*_*before*_ is estimated by 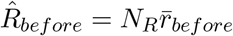 and *R*_*after*_ is estimated by 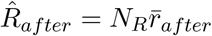. The estimators of the variance are 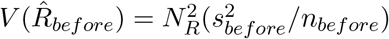 and 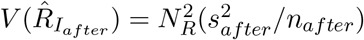 respectively. The time duration between two estimates of *R*_*I*_ before and after the project is implemented, thus defines the time scale of the impact.

This formalization indicates that the sampling unit, *S*_*R*_, does not change from the baseline to the survey. The impact variable is *δ* = |*μ*_*before*_ − *μ*_*after*_|, where *μ* is the average of *r*_*i*_ in all possible samples of *R*_*I*_, before or after project implementation. The inference context consists of formulating a statistic that allows defining the receptor sampling plans in such a way that the estimated impact will be greater than a detection limit (which is always strictly greater than zero). In (Coston-Guarini et al., 2017), a statistic *t* was introduced:

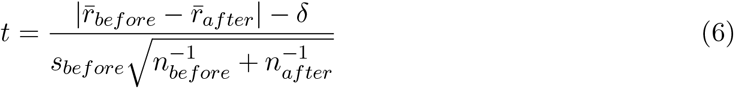

Under the *H*_0_ hypothesis (absence of impact), *δ* = 0, and the statistic *t* follows a Student’s law, with a degree of freedom *ν* = *n*_*before*_ + *n*_*after*_ − 2. Introducing critical values *t*_*α,ν*_ and *t*_*β,ν*_ (corresponding to a first type and second type error, respectively, and which are both fixed *a priori*) the minimum impact inference system is then defined by:

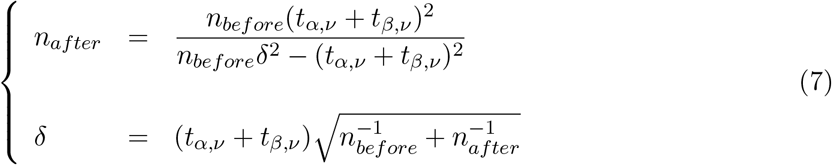

Only *n*_*before*_ is fixed and *n*_*after*_ is determined according to the impact amplitude *δ* and the precision that is required for the EIA study. The *δ* is therefore a minimum impact value allowed by the sampling characteristics. The minimum detection value is 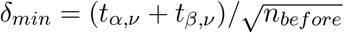 is always positive, showing that assessing an impact is not to decide whether or not an impact exists but to dimension the baseline and survey of the receptor in order to ensure that the impact can be characterized. This inference context requires that the variance remains homogeneous before and after the impact. Hence the ratio must remain close to 1.

### 3.5 eDNA estimates in the context of impact assessment

Instead of observing the receptor presence, the environment surrounding the receptor is sampled in which a quantity of eDNA shed by individuals of the targeted species, *D*_*t*_ is supposed to be present. The challenge for modelling is that the results of different methods of eDNA analysis are not yet consistently quantitative in terms of estimating biomass from sequence data (e.g. review by Rourke et al. (2021) for fisheries). However, promising results from a test of metabarcoding against independent biomass and biodiversity estimates of a marine zooplankton community Ershova et al. (2021), suggest that for some groups this may be resolved in the near future. Nonetheless, once the sample (from the environment) is collected, it is processed to detect the DNA fragments that identify the species. Therefore detection of the species is evaluated in terms of the probability of the presence - or absence - of the species’ DNA being in that sample. The complication with these probabilities is that results can be true, or false positives or false negatives can be obtained (Guillera-Arroita et al., 2017).

For an EIA, it thus becomes necessary to assess the effects of the eDNA protocol on the final outcome. To do this we have modeled a simplified protocol, following Furlan et al. (2016):

- Extraction of DNA from the environment sample. In the present study, it was assumed that the DNA extraction was performed by filtering an aqueous environment. The efficiency of filtering was assumed to be equal to 1.0, in such a way that the number of DNA fragments retained by the filter is equal to the number in the sample.
- Resuspension and homogenization. The DNA fragments retained by the filter were transferred in a liquid solution (volume *V*_*e*_) from which are taken *k* aliquots (each with a volume of *V*_*a*_). The efficiency of resuspension of retained DNA fragments was assumed to be equal to 1.0. In addition, it was assumed that the distribution of the among all possible aliquots is random and hence followed a Poisson distribution law:

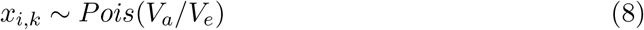
- Amplification and detection of DNA species. Instead of simulating the process in detail, the probability of the amplification success in each of the assays was represented as a Weibull cumulative density function (CDF) of the number of DNA fragments present in the aliquots’ volumes (Furlan et al., 2016):

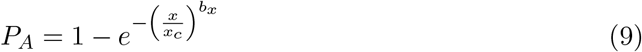

where *x*_*c*_ (in number of DNA fragments per aliquot), and *b*_*x*_ (dimensionless) are two parameters of the Weibull CDF.

It was assumed that false negatives are primarily due to the sensitivity of the amplification and detection method. Therefore *p*_01_ is quantified by 1 − *P*_*A*_. With respect to false positives, each of the samples were replicated k times. The resulting number of positive detections must be compared with the probability distribution to detect 1, 2, 3 … times the species over *k* replicates (Ficetola et al., 2016). The function that describes the probability distribution to detect *y* presences in a series of *k* detection trials, when *p*_10_ (the probability to generate a false positive) is known:

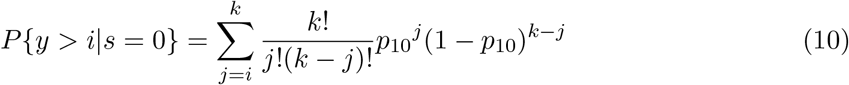

The species occurrence is determined for a number of positive detections over the k replicates that exceed a value for which *P* {*y > i*|*s* = 0} = 0.05. Hence the decision is taken with a first type error equal to 0.05. In addition, according to (Lahoz-Monfort et al., 2016), the probability to detect a species when *y* positive results are found over the *k* replicates is determined by:

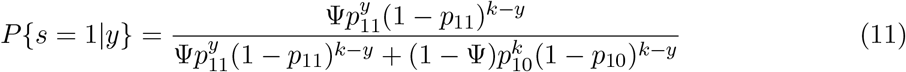

where Ψ is the”site occupancy” probability (the probability that a sample contains eDNA; Lahoz-Monfort et al. (2016)) and *p*_11_ the probability to detect a particular species when eDNA is present in the sample.

This raises several issues linked with sample size:

- if the size of the sample volume is too small regarding the distribution of eDNA in the environment, then many volumes may not contain any DNA fragments of the targeted species. The occupancy probability is then lower than 1.
- if the size of the sample volume is too small regarding the detection of eDNA in the sample (the concentration of eDNA is lower than a detection threshold), and even if the occupancy probability is equal to 1, then the final probability that the species will be assessed as present will be lower than 1.

For the purpose of our simulations in this article, both the probability to generate a false positive and the sample size are known. The replication matrix was calculated, as well as *P* {*y > i*|*s* = 0} and *P* {*s* = 1|*y*} (Ficetola et al., 2016). From *P* {*y > i*|*s* = 0} and the replication matrix, it is decided with a first type error equal to 0.05, that a DNA molecule of the species *s* is present or absent in each of the environment samples. This decision is then compared with the actual (*i*.*e*. simulated) value.

### 3.6 Scenarios to simulate impact

Three sets of scenarios were constructed to simulate the impact of a project implementation on a receptor species. All constants used in the simulation are provided in Table 1 and the scenario conditions are given in Table 2.

**Table 2.** The impact scenarios simulated for a single receptor species using the framework.

| Receptor mobility | Scenario ID | $\phi_R$ | $g_0/g_1$ | $m_1/m_0$ |
| --- | --- | --- | --- | --- |
| Immobile | 1.A | 0.00 | 1 | 1 |
|  | 1.B | 0.00 | 2 | 1 |
|  | 1.C | 0.00 | 1 | 2 |
|  | 1.D | 0.00 | 2 | 2 |
| Passive movement | 2.A | 0.01 | 1 | 1 |
|  | 2.B | 0.01 | 2 | 1 |
|  | 2.C | 0.01 | 1 | 2 |
|  | 2.D | 0.01 | 2 | 2 |
| Active swimmer | 3.A | 0.10 | 1 | 1 |
|  | 3.B | 0.10 | 2 | 1 |
|  | 3.C | 0.10 | 1 | 2 |
|  | 3.D | 0.10 | 2 | 2 |

Notes: The impact scenarios specify receptor mobility ( $\phi_R = 0$ , null; $\phi_R = 0.01$ , passive; or $\phi_R = 0.10$ , active), relative growth ( $g$ ) and relative mortality ( $m$ ). The number of samples collected before and after project implementation were set to 30. The subscript 0 or 1 indicates before project installation, (*e.g.* a baseline) or after impact (*e.g.* when project operation has begun).

In the scenarios, the receptor species is defined as a group of small (*<* 20 cm long) animals, distributed randomly over a distribution area, which includes the project area. The individuals living within the project area are submitted to an impact. In the absence of an impact, before project implementation, the distribution of the receptor species at equilibrium 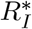 was assumed to be equal, within and outside of the project area, on average. This leads to the description: *R*_*E*_ = *R*_*max*_(*g* − *m*)*/g*. The eDNA distribution was also assumed to be equal, on average, over the distribution area, leading to : 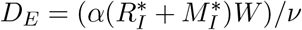.

The receptor species is allowed three different capacities of movement: no movement capacity, passive movement and active movement. As DNA molecules are assumed to disperse passively in the water column, the movement capacities were converted to *ϕ*_*R*_ = 0 for no movement, *ϕ*_*R*_ = *ϕ*_*D*_ for passive dispersion and *ϕ*_*R*_ *> phi*_*D*_ for active movement. In the latter case, movement is still conceived of as dispersive, but as the result of a random walk of all individuals at a faster speed than if they were dispersed passively, only by environmental mixing.

The project impact was assumed to affect the physiological performance of the receptor, represented by dividing the growth rate *g* (hence, *a*_*g*_) and multiplying the mortality rate *m* (hence, *a*_*m*_) by a factor of 2, successively and together. The impact simulation plan therefore crossed three conditions for the movement capacities of the receptors and three conditions for the impact. We focused on mobility in our scenarios because this emphasized exchanges of individuals in and out of the project area. However, different criteria may be more relevant in other impact studies.

### 3.7 Coding

The model and F-statistic test were programmed using the open source numerical computation tool, SciLab (v. 6.1.1, https://www.scilab.org/). Results of simulations were analyzed and plotted in both SciLab and Datagraph (v. 5.1, Adalsteinsson 2022).

## 4 Results

We begin by summarizing the results of the analysis of the mathematical properties of the model. Then the results for the equilibrium states of both the receptor and eDNA and their stability, and the time required to reach new equilibrium values after a disturbance are presented. Finally, the results of the simulation framework (Figure 1) tested with the twelve different plausible impact scenarios (Table 2) are shown.

### 4.1 Stability of the receptor

The observation of a sample of *R*_*I*_ leads to the estimate of 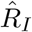 and its uncertainty. Nonetheless, for assessing an impact, the receptor must not be in a transitory phase at the time scale of the impact. It should be close to a stable steady state (*i*.*e*. the average should not drift and the variability at small scale should be assimilated to a random noise), before and after project implementation.

To ensure this condition, the stable equilibrium values of the Equations 1 with 2, and 3, were determined:

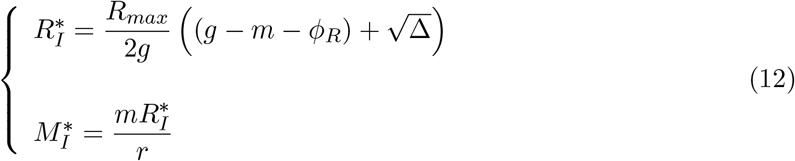

where Δ = 4(*g/R*_*max*_)*ϕ*_*R*_*R*_*E*_ + (*g* − *m* − *ϕ*_*R*_)^2^ is a discriminant.

It is worth noting that both Equations 1 with 2 have two steady state solutions; only the positive one is retained and is determined by the property that 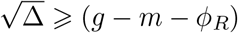. This solution emphasizes that when the exchange rate *ϕ*_*R*_ is greater than zero, the receptor quantity is partially conditioned by the individuals located outside the impacted zone (here, equivalent to the project area) that would be considered as non-impacted by the project under EIA. When the exchange rate is null, the stable equilibrium is a unique solution equal to 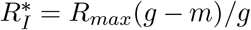 and positive only if *g* is greater or equal than *m, i*.*e*. if the growth rate is greater or equal than the mortality rate. As an equilibrium can be reached before and after impact, changes in parameters {*g, m, ϕ*_*R*_} induce a change in 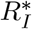. Hence, the impact is quantified by the difference between the two stable steady-state values.

### 4.2 Variability of eDNA at receptor equilibrium

At the equilibrium of *R*_*I*_ and *M*_*I*_, it is possible to integrate analytically Equation 5. The dynamic of *D*(*t*) is then formulated as:

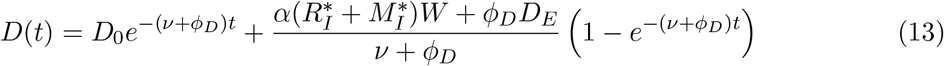

The equilibrium for *D*(*t*) is then 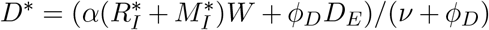 and is reached when *t* tends to +∞. In practice, this solution demonstrates the importance of quantifying the time delay between the states of the receptor and the corresponding eDNA values. This becomes important primarily after project implementation, when transitory phases may lead to impacted steady states from non impacted ones.

The time to reach a value close to a steady-state is equal to:

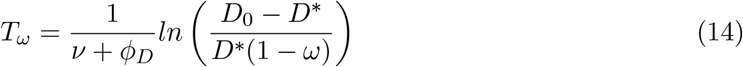

where *ω* is a proximity coefficient close to 1 (*ω* ⩾ 1 if *D*_0_ > *D*^*^, and *ω* ⩽ 1 if *D*_0_ < *D*^*^). If *ω* = 1 then *T*_*ω*_ is equal to +∞. In practice, *ω* must be set equal to the small scale random variability of the state variable *D*.

Since *D*^*^ depends on *ϕ*_*D*_, this result also emphasizes the influence of the spatial dimension on the difference between the receptor(s) state(s) and its shed DNA. Generally, in the project area, even if it is possible to calculate a (simplified) steady-state solution, only *D* can be expressed (and sampled), while the actual quantity of interest, *D*_*I*_ that corresponds to the impacted receptor *R*_*I*_, would remain non-observable.

### 4.3 Simulation of an impact

Since the project impact is on the ecophysiological properties of the species, it is assumed to be detectable through a decrease in the receptor abundance. All sample density averages, before and after project implementation, were greater than 15. The variance ratio, before and after project implementation, remained close to 1. These two conditions validated the approximation of a Poisson distribution of the estimator by a Normal distribution and the impact estimates to be performed.

Apart from the control simulations (*i*.*e*. no changes of parameters were simulated), a sampling effort of 30 samples taken before and after project implementation was enough to detect, in all cases (null, passive or active mobility and growth divided and/or mortality multiplied by 2), an impact on the population state. The effect simulated was indeed large enough to trigger a difference greater than the minimum detection value *δ* (Equation 7). In most cases, only three samples would have been enough to be able to detect a significant decrease. In this case, the minimum detected change, which only depends on the spatial distribution of individuals and number of samples before and after project implementation, is *δ* = 0.69 *ind*.*m*^−1^.

The only two exceptions are for the case of a receptor species with the capacity for active displacement. First, when growth was divided by two and mortality was not affected (Table 2, 3.B), seven samples are required to monitor a change in average greater than *δ* = 0.34 *ind*.*m*^−1^). Second, when growth was not affected but mortality was multiplied by two (Table 2, 3.C), then four samples are required to monitor a change in average greater than *δ* = 0.51 *ind*.*m*^−1^).

**Table 3.**
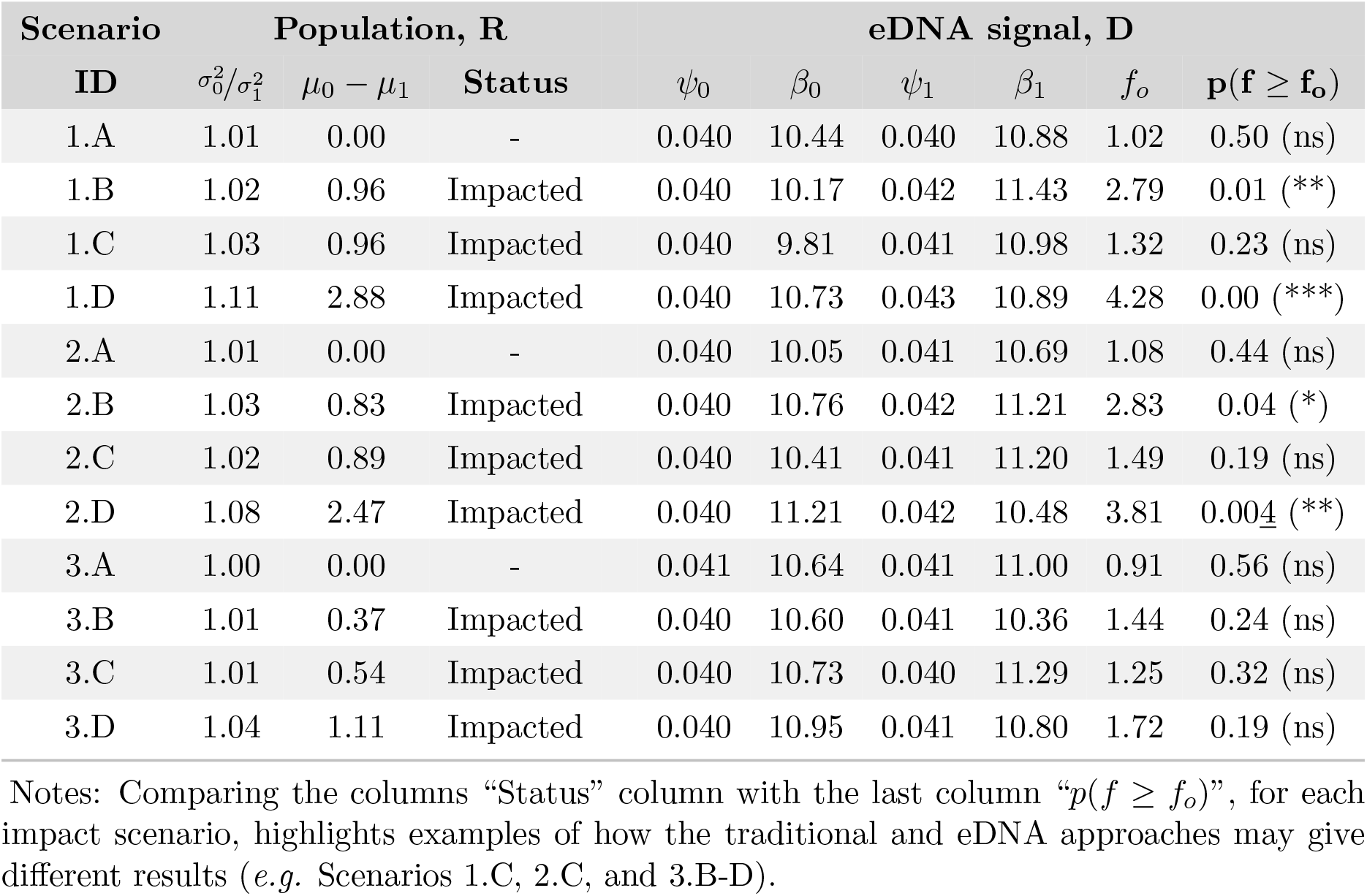
Results of the 12 impact scenarios simulated using the population (R) or eDNA signal (D) of a single receptor species.

For eDNA in all twelve scenarios, a successive series of 200 samples with sampling units (*S*_*D*_) that varied from 0.004 to 0.060*m*^−3^ every 0.002*m*^−3^ were simulated in order to detect the presence of the species from eDNA analysis (Figure 2). The true presence was assessed using Equation 10 with *p*_10_ = 0.1 and *y >* 4 over *k* = 8 replicate analyses to ensure that the first type error was lower than 0.05 (see Equation 10).

**Figure 2:**
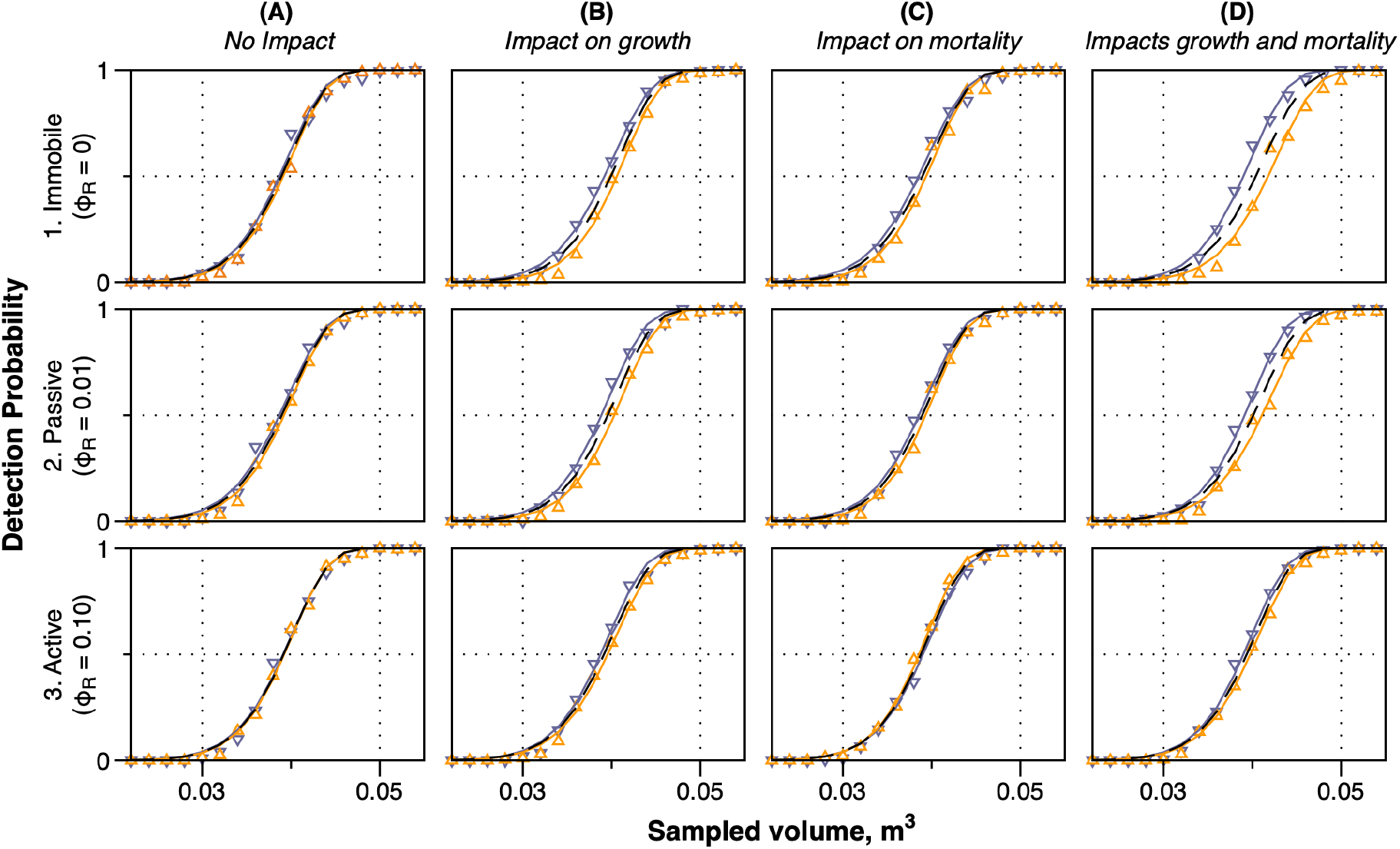
The eDNA detection curves of a receptor species simulated for all 12 scenarios (Table 2) with 200 samples each. Three types of samplings are treated: baseline (purple), after project implementation (orange) and the statistical null case (black, dashed line). In ten cases, the receptor species would be detected for a sample volume between 40 and 50 L. Only two scenarios (1.D and 2.D, where impact on an immobile or passive species is expected to disturb both growth and mortality) could require larger sample volumes. To make visible the small differences in slopes of some cases, the volume range plotted was limited to 0.022 - 0.055 m^3^.

The detection function first checked that the occupancy Ψ was equal to 1 in all sample sets. Then the results of the occupancy model, *P* {*s* = 1|*y*} (Equation 11) were calculated. They exhibit strong fluctuations for small sampling volume detection, even if the occupancy Ψ was constantly equal to 1, and Ψ, *p*_10_, *p*_11_, *k* and *y* were known and not subject to estimation. Then the detection curve was calculated which links the presence probability to the sampling volume. To do this, detection results are filtered into false positives and false negatives, at a threshold of 0.05 (Type I risk, Equation 10). This retains a minimum of four positive detections for a series of eight replicate analyses of the aliquot; these points are plotted on Figure 2.

The resulting series of detection probabilities as a function of the sampled volume were fit with a Weibull cumulative distribution function (CDF):

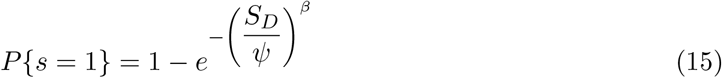

where *β* is the shape parameter and *ψ* is the scale parameter, expressed here as a unit of volume.

For each of the tested combinations {*ϕ*_*D*_, *g, m*}, three sets of best parameter estimates {*ψ*_0_, *β*_0_}, {*ψ*_1_, *β*_1_}, before and after project implementation respectively, and {*ψ*_*T*_, *β*_*T*_ } for all data pairs pooled together, were estimated by optimization. The optimization of such a distribution should use a maximum likelihood estimator. However, the convergence of the estimator depends on *β*, and the estimate of this parameter requires a numerical evaluation. Therefore it was decided to estimate both parameters at once, minimizing the residuals squared sums between observations and predictions (*RSS*_0_, *RSS*_1_ and *RSS*_*T*_, respectively; Equation 15) using a direct search algorithm (Simplex method of Nelder & Mead (1965)); estimators developed by Menon (1963) were used to initiate the simplex algorithm.

Finally, in order to decide if there is a significant difference between the two curves, before and after project implementation (and also between the two corresponding sets of parameters) an

F-like statistic, *f*, was introduced:

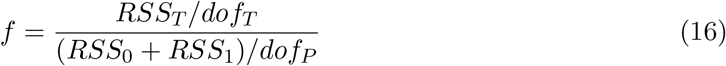

where *dof*_*T*_ and *dof*_*P*_ are degrees of freedom *dof*_*T*_ = *N* − 2 and *dof*_*P*_ = *N* − 4, *N* been the total number of observations, with two parameters to estimate, once for *RSS*_*T*_ and twice for *RSS*_0_ + *RSS*_1_.

The distribution of the F-like statistics was calculated analytically because the underlying distribution models for the residuals cannot be assimilated to a centered and reduced normal distribution. Thus the residuals squared sum (*RSS*) cannot be assumed to follow *χ*^2^ distributions. Therefore, empirical distributions were estimated for each simulation scenario, by bootstrapping 300 times, the centered residuals corresponding to *RSS*_*T*_. This means the null hypothesis states that filling one model with all data is equivalent to fitting two models with two separate data sets. The bootstrap procedure recreates 300 pseudo data vectors {*ψ*_0_, *β*_0_ *ψ*_1_, *β*_1_ *ψ*_*T*_, *β*_*T*_ }, re-estimating 300 times *RSS*_0_, *RSS*_1_ and *RSS*_*T*_ (Efron & Tibshiraini, 1993) and calculates the probability that *f* exceeds *f*_*o*_, the nominal value of *f* calculated using the best estimates.

The results of these comparisons for all scenarios are summarized in Table 3. The situation of no impact was tested for each type of mobility scenario, leading logically to the non-rejection of the null hypothesis. For the scenarios with an immobile receptor, a significant difference (with a first type error lower than or equal to 0.01) between the two detection curves, before and after, was assessed for both scenarios in which growth rate was divided by two (scenarios 1.B, 1.D). However, no significant difference was assessed when only mortality was multiplied by two (growth remained unchanged, 1.C), despite the fact that an impact was assessed for the receptor (Table 3). A similar set of results was obtained for scenarios corresponding to a passive movement of the receptor (1.A-D), which is expected to be identical to the dispersion of DNA fragments in their environment. However, the rejection of the null hypothesis required to take a higher first type risk (0.05 instead of 0.01). Finally, for scenarios with active movement (*i*.*e*. random walk faster that passive dispersion, 3.A-D), all results, whatever the combination of ecophysiological effects was, show no significant differences between the two curves, before and after the project implementation and even if impact was assessed on the receptor in all cases.

Lastly, we can also explore the range of sample volumes required to detect impacts. By taking into account the parameters we have set in our simulation framework, mainly accounting for an average receptor density of 31 *ind*.*m*^−2^ in a project area covering 1.0.10^5^*m*^2^, with a water height of 3 *m*, we can find that the cut-off volume for the detection is between 40 − 50 liters, which is much higher than volumes typically sampled. It can even increase to 325 liters, with a shape parameter of *β* = 10.5, when the lowest shedding rate and highest decay rate (in their range of variability) are applied (Table 1). Even in the case where the highest shedding rate and lowest decay rate are applied, the scale volume is still approximately 6 liters with a shape parameter of *β* = 12.

## 5 Discussion

Our study combined a classic inference framework that assesses impact on a receptor species with a heuristic framework in order to explore how species detection using eDNA analysis could be used within EIA. To support and encourage experimentation with this approach, a deterministic model was also developed to simulate observations for both the abundance of the receptor species and the quantity of DNA released in the environment, before and after project implementation. The objective of the framework presented is to provide a complete representation of the dynamics of both the receptor and its shed eDNA, such that states can be compared with estimates performed when simulating sampling plans (Figure 1).

### 5.1 Summary, advantages and limitations of a deterministic framework

The deterministic model was designed without spatial dimension and simulates the global dynamics of the state variables within a project area. The spatial distributions were calculated in a second step defining the statistical context for the study. Random spatial distributions were chosen to be minimal, in such a way that the only sampling plan that can be applied is a simple random sampling. With this approach, the variances (which define the uncertainties of the estimators) have the same amplitude as the averages. Despite these choices, the simulated effects ensured that impacts can be assessed without ambiguity, and that the differences in the receptor state estimates were significant even with a small sampling effort.

The framework is conceived of as a field-to-laboratory numerical pipeline system. This means we represent a detection of the species that depends not only on the presence of DNA in the environmental samples, but also on the subsequent DNA amplification and sequence identification. Three steps in eDNA analysis are typically described by three successive probabilities (McClenaghan et al., 2020):

- *occupancy* estimated by the probability that eDNA is present in a site, given that the species is present;
- *capturability* estimated by the probability that eDNA is extracted from a sample, given that eDNA is present in the site; and
- *detectability* estimated by the probability that the species is detected, given that eDNA was extracted from the sample.

This probabilistic context was simplified by ensuring that the eDNA occupancy was equal to 1, even when the impacted receptor abundances decreased, and that the extraction efficiency was equal to 1, as well. In this context, the probability to detect the species by eDNA analysis when the species is present in the sample (which is *Occupancy* × *Capturability* × *Detectability*) was condensed to detectability only, in our approach. This emphasizes that an impact is not supposed to be assessed by eDNA presence detection (as long as there is no complete disappearance or new introduction of the species after project implementation), and, conversely, that looking for a criteria to detect a change must be based on the detectability, because, otherwise all other probabilities are assumed to be equal.

The steps of eDNA analysis (amplification, sequencing and identification of the sequence) were simulated as described in (Furlan et al., 2016) for a single species specific assay. The protocol was simplified and does not account for error propagation and loss of efficiency at the different simulated steps. The outcome is a species detection probability, that varies between 0 (if the species cannot be detected from samples) and 1, which means there is a certainty to detect the species from all samples analyzed. This property was used to build detection curves, linking detection probability *vs*. sampling volume curves before and after project implementation, to test their variability and infer eventual differences. The detection curves presented can be thought of as a cumulative probability distribution, where the sampling volume is the random variable that accumulates progressively the probabilities as it increases.

### 5.2 Sampling plans and preliminary tests are strongly recommended

For a ‘traditional’ organism-oriented approach to mapping the distribution of a receptor (or a targeted species), a sampling plan can only be optimized for the population for which a state has been defined. In contrast, an eDNA-oriented sampling plan will use a sampling unit *S*_*D*_ and total number of samples, *m*, which could require many more samples than the traditional approach. In the modeling presented, we emphasized this difference by re-scaling the eDNA sampling domain regarding the receptor. Hence, in the absence of any prior knowledge, the two applicable sampling plans would be: simple random sampling (that is, if no structure can be hypothesized) or systematic sampling (if a continuous structure can be hypothesized). However, the location, direction, intensity and extent of a molecular gradient(s) is(are) difficult to determine without understanding both hydrodynamics and the dynamics of eDNA in the environment. Here, we assumed that the eDNA distribution was random. Nonetheless, the problem of spatial and temporal structures should be explored further, since unexplainable differences between samples taken only a week apart have been reported (Beentjes et al., 2019).

Finally, for the sake of simplicity, a sampling was assumed to be performed with replacement in the simulations. This means that there is very little difference between the ‘with’ and ‘without’ replacement condition, as long as the number of samples is very small with respect to the total number of samples possible. The organism-oriented approach, if it uses a sample number of 100, then the number of realized over total samples would be equal to 0.00007. For an eDNA sampling plan, even with the largest volume filtered used of 0.250*m*^3^ and 200 samples collected, the number of realized over the total samples would be equal to 0.00008.

### 5.3 Assumptions and limitations

The system formulated for this study was assumed to work as a meta-(eco)system (Loreau et al., 2003). The local, demographic dynamics of the receptor is supposed to be different within the project area (where the receptor is subject to impact) from outside the area. Regional exchange is represented by a mixing process, described by a mixing rate between the compartments inside and outside of the project area. This description is only valid if there is no continuous gradient. Artificially, this was ensured by assuming that the receptor was distributed randomly inside and outside of the project area. When a continuous structure exists, the structure (*e*.*g*. spatial covariance) must be quantified to an advantage in order to increase the precision on estimates. In the meta-(eco)system, horizontal mixing was kept low, and thus exchanges outside of the project area were limited, so that the local dynamics of the receptor species within the impacted area predominated over regional dynamics. This situation allows impacts to become significant. In the absence of impact, it was assumed that receptor abundances within and outside of the project area were equal; however, the calculation of *R*_*E*_ only depended on demographic parameters {*R*_*max*_, *g, m*}.

A reactive-transport type model allows identification of the horizontal structure induced by water mass movement. These have been used with success in a small number of eDNA studies. For example, Sansom & Sassoubre (2017) studied the spatio-temporal variability of eDNA shed by freshwater mussels (*Lampsilis siliquoidea*) within a river system. They supported their observation (data collected in both experimental and *in situ* conditions) with simulations from a reactive-transport model. Their modeling identified the spatial gradient of eDNA concentration from the mussel bed, at equilibrium. More generally, Andruszkiewicz et al. (2019) promoted the use of process-oriented models to understand the spatio-temporal distribution of eDNA, attempting to reconstruct, from observations, the origin in space and time of eDNA shed by anchovies. Also, Ellis et al. (2021) have developed a similar model and compared simulations with observations to quantify the spatial range for the detection of eDNA shed by two sources: a kelp and a sea star species. They found very similar results regarding the order of magnitude of both the horizontal extent, and the duration to reach a detection limit from a shedding time. Finally, Allan et al. (2021) used a similar modeling approach but at a much more refined scale to investigate the vertical distribution of the eDNA in relation to the source of shedding. They studied the relative importance of eDNA transport and organism movements, finding that the effect of eDNA vertical transport was negligible for the potential nycthemeral migration behaviors of planktonic organisms.

Modeling is foreseen as a means to understand the dynamics of DNA released in the environment, however, the primary challenge in designing and using models to quantify the dynamics of eDNA is their parameterization. In our approach, only three processes were accounted for: transport, shedding and decay. They were assumed to be generic phenomenological processes that can be represented by a minimal number of parameters. Each of these processes encompasses several mechanisms having a different importance, according to the study case (Barnes & Turner, 2015):

- Transport encompasses not only advection and dispersion (isotropic or anisotropic in three dimensions), but also settling and resuspension, connecting the dynamics of eDNA between different environments and governed by other mechanisms, such as biomixing and bioturbation. Secondary processes must be important like transport of eDNA fragments by particles or organisms.
- Shedding, which is a release of eDNA fragments, can be performed by living and dead organisms. It has important implications for EIA studies, only concerned by assessing effects on living organisms. In addition, shedding can be passive and/or active, hence including behavioral components in their kinetics. As for transport, a set of secondary mechanisms must be invoked and accounted for which are linked with trophic mechanisms.
- Decay, designates a global loss of eDNA once shed in the environment, and can be subject to very different interpretations according to the context of studies. The primary set of mechanisms are encompassed under degradation, which can be biological, chemical or physical. However, losses can also be due to mechanisms that prevent eDNA to be sampled, detected or identified when it changes in composition or other physico-chemcial characteristics.

To the complexity of the processes, due to the multiplicity of mechanisms, is added the effect of abiotic environmental conditions (*e*.*g*. light, temperature, salinity, pH, …), which can be relevant for EIA studies. Abiotic factors also affect eDNA according to their state (*e*.*g*. intramembranous or extramembranous, free or bound), configuration or conformation, when it is exposed (Barnes et al., 2014). When reviewing existing published studies (Hinz et al., 2022), we found that there is very little quantitative information about processes that govern eDNA dynamics at such a level of complexity. Frequently, global shedding and decay rates are estimated. Estimates exhibit not only a large inter-specific and environmental variability, but also a large random, unexplained, variability in controlled conditions experiments (Sansom & Sassoubre, 2017). This has led many authors to use the more general and fuzzy concept of the “persistence of eDNA” in the environment (Barnes et al., 2014; Collins et al., 2018; Salter, 2018). The term ‘persistence’ is ambiguous because it encompass a notion of presence (a balance between shedding and decay), and a perception of a duration, linked with the idea that the molecules progressively disappear or escape from detection by decay, dilution, burial, or transport.

In our approach, we have chosen to stay as generic as possible, but in order to perform simulations, we were obliged to set values for all parameters of the model. Some parameter values estimates with different units are available in the literature (*e*.*g*. Allan et al. 2021). We chose to use the average values determined by (Sansom & Sassoubre, 2017) because these were obtained for densities of organisms close to the densities in our system. They were acquired in experimental conditions and did not exhibit variations as a function of the organisms’ density. However, we were not able to consider that shedding and decay rates can vary before and after project implementation. Indeed, (Stewart, 2019) pointed out not only that both rates can vary according to the abiotic environmental conditions but also, concerning shedding rates, according to other biotic factors, including stressors. Our approach therefore remains minimal, and a much larger complexity could be expected with more associated sources of variability in concrete cases.

Another source of complexity is inherent to the analytical process used to detect species from sampling eDNA in the environment. Barnes & Turner (2015) underscored the technical challenges associated to the study of the different processes governing eDNA dynamics, distinguishing the technical issues *per se* (for example, linked to the efficiency of the different methods) from the potential sources of errors (for example, due to contamination or sequencing mistakes). The most striking issue is nonetheless the partial disconnection between the sampling step and the analyses. This partial disconnection occurs when eDNA is extracted from the sample, eluted and then re-sampled for replications. Simulating these analytic protocols requires knowing the quantities that are needed for the evaluation, which are the extraction (filtration) and elutant volumes and the aliquot (*i*.*e*. replicate) volume. It also requires knowing the amplification efficiency and analytical limits of detection. The resulting probability to detect a species is conditioned by the ratio between the elutant and the aliquot volumes (Furlan et al., 2016). Surprisingly, this ratio seems to be imposed by “kits” used to extract and amplify DNA, without considering the particular conditions imposed by the studied case.

### 5.4 On the use of eDNA to detect impact

For EIA, environmental DNA analysis is is well positioned to work as a complementary tool on many tasks, like detecting: invasive species, the continual presence of endangered species, or enlarging the spectrum of receptors of potential impacts. However, eDNA was conceived of as an exploratory (descriptive) method, rather than an inferential framework compatible with Environmental Impact Assessment. Although the technique of qPCR provides an estimate of the initial eDNA quantity based on the number of cycles, threshold and calibration curves, the propagation of errors and decrease of efficiency at different steps of the process also generate a large uncertainty and potential bias on the final result. In addition, impact assessment concerns a concept of the “receptor” which is typically, a group of organisms or a sub-population living within the project area. The receptor can also be an assemblage of individuals belonging to different species attached to a specific habitat within the project area. Combined with complex environmental dynamics, this creates many challenges for the interpretation of eDNA signals, if adequate preliminary testing and suitable sample plans have not been prepared.

In addition, the challenge of quantifying impact with eDNA concerns more than analysis methods and protocols. It requires to modify the approach to impact assessment itself, because comparisons of series of detection curves, like we have generated here, substitute for decision theory. Decision theory is not about tests to see if there is a change, but instead it is about understanding what needs to be done to characterize a predicted change Coston-Guarini et al. (2017). Using a detection curve lifts ambiguities that the eDNA data can contain.

However, environmental DNA will become important for EIA studies if it succeeds to assess changes on receptors under the spatial and temporal constraints of a Project Area and Project Lifecycle. There are three areas of concern:

- When samples of an environment contain a mix between the eDNA shed by living organisms within and outside of the project area, and shed by dead organisms, all from the same species, it is difficult to identify the fraction of eDNA shed by the receptor. This may be resolved with eRNA instead Yates (2021).
- The distribution of eDNA among samples can be truncated when sampling units are too small. Many zero occurrences in samples can prevent the presence of the species to be assessed, despite of the fact that the receptor is present in the project area, and hence occupancy is equal to 1. There is no acceptable principle to optimize the sampling plan for eDNA *a priori*, because it depends on many unknown components, starting by the unknown state of the receptor population(s).
- A fraction of the eDNA concentration distribution can end up being below the detection probability, generating a large quantity of false negatives in samples. This can happen despite the fact that the eDNA of the targeted species is present in the extraction eluent and replicates.

In our heuristic approach, to construct curves that can be used in EIA, we exploited the probabilistic property of eDNA analysis: the detection probability theoretically increases asymptotically from 0 to 1 when the sample volume increases (Darling & Mahon, 2011). By testing a range of sample volumes in our sampling plan, a curve linking the detection probability (probability to detect the true presence of species) with the sampling volume could be constructed. A Weibull CDF function was fitted to the data and F-like statistics compare the two curves (the baseline and after project start). This comparison tests if a difference is significant when an impact has been assessed on the receptor.

Comparing the detection curves before and after the simulated project implementations (Table 3 and Figure 2) allowed for the assessment of significant differences in a limited number of cases: when the receptor growth was affected and when the mobility of the receptor was limited, null or passive. Under a passive movement scenario, the local dynamics would not interfere with spatial exchanges of organisms, accounting for the characteristic that eDNA is dispersed passively. For the case of active dispersion, whatever the combination of effects was, no significant difference between detection curves before and after impact were found. This suggests that if a receptor species is an active swimmer, eDNA would not be suitable for impact forecasting. Nonetheless these results cannot be generalized and we caution the reader that we are working within the construct of a thought experiment. Results remain completely dependent on the conditions of the simulation, the physical dimensions of the system, the biological characteristics of the species, the impact characteristics (*i*.*e*. the simulated effect), the processes governing the eDNA dynamics and the eDNA analytical protocol.

Our results do demonstrate, however, that an absence of significant difference between detection curves before and after project implementation cannot be interpreted as an absence of impact. *A contrario*, a significant difference indicates a change in eDNA concentration and spatial distribution. However, it should be interpreted with caution regarding an impact on the receptor without considering carefully what other conditions could have changed.

In environmental and ecological sciences, statistics alone cannot compensate for sub-optimal sampling. Environmental DNA, shows that without to understand underlying mechanisms, processes, dynamics and structures, there is very little chance that the state of the receptor species can be characterized. Kelly et al. (2016) have found counter-intuitive results of diversity pattern while sampling eDNA in a urbanization gradient, experiencing difficulties to interpret their results. Indeed, many authors (*e*.*g*. Barnes & Turner 2015) working on eDNA applications have appealed for a better understanding of the “fate” of eDNA, from the moment it is released to the moment it is sampled and analyzed. Several authors (*e*.*g*. Andruszkiewicz et al. 2019; Allan et al. 2021; Carraro et al. 2020) have promoted an approach that combines statistic estimates and mathematical modeling of the dynamics of eDNA concentrations in aquatic systems to explain distribution patterns and contribute to sampling plan optimization. They suggest that precise parameter estimates are required, and this is a position we also agree with.

## 6 Conclusions

By using a comprehensive field-to-laboratory simulation, we were able to infer differences in eDNA detection probabilities concomitant to the existence of an impact. The development of the tool, the construction of the detection curves (linking the detection probability to environmental sample volumes), and the formalization of a statistic for a non linear model (supported by a Monte-Carlo method to estimate their null distribution) offers new perspectives to examine the roles that eDNA may have within EIA studies.

Some caution should be exercised with EIA. Two major limitations have been identified. First this approach is expected to require additional effort to be fully implemented. This is because several sets of samples over a large range of sample volumes would be needed, both before and after project implementation. Our approach can be expected to multiply both the effort and costs that eDNA is supposed to decrease. Second, there is a strong need to develop precise descriptions of the dynamics of eDNA reactivity under diverse conditions. Rigorously pre-testing the modalities of the analytical protocol under study specific conditions is recommended. All of this would entail additional effort, which is not usually part of the scope of Environmental Impact Assessment activities.

Nonetheless, it is hoped that this work will inspire further work to develop a systems approach to eDNA with numerical simulations and new statistical inference tools, as well as incite others to develop new applications of eDNA data suitable for environmental impact studies.

## Data Archiving Statement

There are no data in this study.

## A. Supplementary Information

This section contains additional details about the model parameterization that was described in Table 1 of the main text, as well a background information on sampling volume choices used in aquatic and marine systems compiled from a literature survey of more than 600 articles, discussed in an earlier article (Hinz et al., 2022).

### A.1 Dimensioning the system in the model

*A* in (m^2^) is the project area, that is the area in which there is an impact to forecast.

*ρ* : A scaling environmental factor : the receptor is defined according to a specific environment (for example, a benthic species or community is expressed in term of quantity on a surface). eDNA is defined according to the environment that contains it. For the benthic species, expressed as a function of a surface, either for sediment or overlying water, eDNA is expressed as a quantity per unit of volume (as it is sampled). *ρ* scales the receptor’s dimension to the eDNA’s dimension (here a surface to a volume, hence multiplying the surface by the water height or the sediment depth in which eDNA can be found, here, in m).

*S*_*R*_ (in m^2^) the Sampling Unit for the receptor

*S*_*D*_ (in m^3^) the Sampling Unit for eDNA. This cannot be optimized so a range of volumes is applied. Sampling Units are fixed values for one sampling: so a range of sampling volumes implies a range of different samplings.

*A* was chosen in such a way that *ρA > A >> Su*_*R*_ and *ρA >> Su*_*D*_. In a practical manner, the choice was guided by computational effort and simulation time.

### A.2 The Receptor

*L* (in mm) is a measure of the organism used in allometric functions. Allometric functions are described as 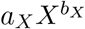. Allometric relationships are dimensional (*e*.*g*. a volume is a cube of a length), and respect the “Energetic Equivalence Rule” (Damuth, 1991).

*a*_*L*_ : (in g.mm^−3^) coefficient for the length to weight relationship, considered for *b*_*L*_=3

*a*_*R*_ : (nb of individuals.g.m^−2^) coefficient for the weight to maximum relationship, considered for *b*_*R*_ = -1

*a*_*g*_ : (g^0.25^.day^−1^) coefficient for the weight to growth relationship, considered for *b*_*g*_ = -0.25

*a*_*m*_ : (g^0.25^.day^−1^) coefficient for the weight to mortality relationship, considered for *b*_*g*_ = -0.25

### A.3 Describing Exchange rates

*ϕ*_*D*_ (in time^−1^) is an eDNA dispersal rate determined arbitrarily but is considered as “passive”, since eDNA does not have any motility.

*ϕ*_*R*_ (in time^−1^) is the receptor dispersal rate determined arbitrarily. The receptor is considered as “passive” or “planktonique” if *ϕ*_*R*_ = *ϕ*_*D*_, null (sedentary) if *ϕ*_*R*_ =0, or “active” (“nectonic”), if *ϕ*_*R*_ *> ϕ*_*D*_. In the latter case, *ϕ*_*R*_ is still a dispersal rate but due to Brownian motion, it is faster than the one triggered by the motion of the water mass.

### A.4 Including eDNA processes

These are all written in Latin characters.

_*αW*_ is the weight-specific shedding rate (DNA fragment shed.h^−1^.g^−1^)

*ν* is the decay rate (h^−1^)

### A.5 Representing the eDNA analysis protocol

*V*_*e*_ : (in ml) Eluent volume: it represents the volume in which eDNA is placed, obtained after a series of manipulations to extract eDNA from the environment sample. It is associated to an overall efficiency (between 0 and 1), here considered to be 1. *V*_*e*_ is usually defined by kit protocols (Furlan et al., 2016).

*V*_*a*_ : (in *μ*l) aliquot volume: it represents the volume taken from Ve, containing a random franction of the sampled and extracted eDNA. It is a replicated volume (see parameter *k* below) submitted to an amplification and detection analysis. Usually defined by kit protocols (Furlan et al., 2016)

*x*_*c*_ : is a scaled number of fragments used in the efficiency curve for the technique used to amplify eDNA fragments. Taken from Furlan et al. (2016).

*b*_*x*_ : is the shape parameter of the efficiency curve (the higher, the sharper). Taken from Furlan et al. (2016).

*p*_10_ : probability to “detect” a species when this one is not present. It depends on the techniques and protocol (but also the skills of the analyst to avoid contamination). It is then fixed here arbitrarily but still low regarding the level of probability to detect a species when present.

*k*: number of replicates/aliquots/assays. It comes after *p*_10_ because, usually, it is necessary to know *p*_10_ to calculate the Bernouilli distribution of positive and/or negative detections, and fix a threshold to decide if the number of positive detections indicates a true presence or not.

### A.6 How is the choice of sample volume presented in the literature?

During a previous review article on eDNA (Hinz et al., 2022), we examined more than 600 publications (n = 641) to understand the reasoning underlying the choice of water sample volumes in eDNA studies.

We were surprised to find that only three review articles dealt explicitly with the question of sampling volume. In general, we found many articles justified their sample volumes by referring to an earlier suggested sample volume, particularly from Ficetola et al. (2008) or Jerde et al. (2011). We also noted that many of the papers examined did not actually mention the volume collected explicitly, leaving the reader to make some assumptions, based on the filtration described or bottle sizes.

Appendix Figure A.1.A shows the distribution of sample volumes compiled from 41 different sources (see sub section A.6.1). Cited sampled water volumes range from 5 mL to 30 L. The smallest volumes are the volume recommended by a particular extraction kit, and the largest volumes are used in a range of aquatic and marine studies, as well as for experimental (aquarium, or pond) studies.

The second plot (Appendix Figure A.1.B) shows the distribution of volumes that have been characterized as “typical”, “normal” or even “standard” for different environments. We could not find any justification for these strong qualifications.

We also remarked that if preliminary testing was done in studies, it has not been mentioned. There indeed appears to be no quantitative evaluation of the volume choices being recommended in the majority of studies.

This implies an important and fundamental oversight in the methodology and reporting practices of eDNA studies. Nearly every eDNA article mentions the basic fact that detection depends on the amount of DNA mass in the sample, and therefore the size of the original sample. However, a volume size cannot be both a reason for a non-detection and not have been subject to preliminary testing. This also undermines the utility of this reporting for future studies, as well as the re-use of published data to: re-analyse trends, establish baselines, perform meta-analyses *etc*..

We therefore join other authors, such as Allan et al. (2021), in strongly recommending that the sampling volumes be evaluated and chosen within the context of eDNA studies with both statistical sampling tools and preliminary tests to determine the precise conditions of the detection curves for the targeted organisms in a particular set of conditions.

**Figure A.1.**
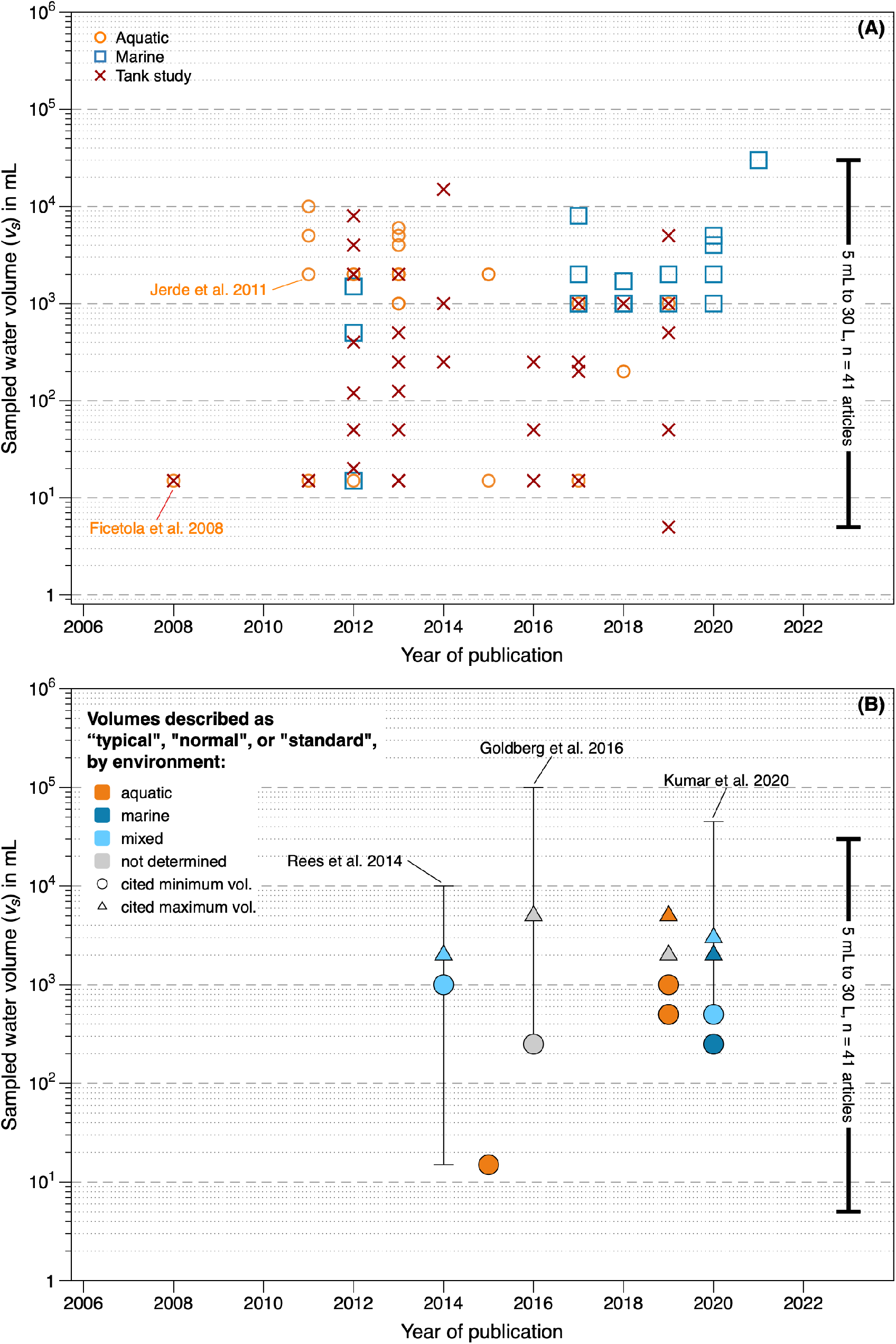
Plot (A) shows the distribution of water sample volumes reported in the eDNA literature, by water type. Plot (B) shows the distribution of sampling volumes described as recommended, by water type.

